# Aging-related hypometabolism in the anterior cingulate cortex mediates the relationship between age vs. executive function but not vs. memory in cognitively intact elders

**DOI:** 10.1101/635219

**Authors:** José V. Pardo, Shantal M. Nyabwari, Joel T. Lee, ADNI

## Abstract

Elucidating the pathophysiology of cognitive decline during aging in those without overt neurodegeneration is a prerequisite to improved diagnosis, prevention, and treatment of cognitive aging. We showed previously the anterior cingulate cortex (ACC) and adjacent medial prefrontal cortex (mPFC) are centers for aging-related metabolic dysfunction that correlate with age-associated cognitive decline in healthy volunteers. Here, we examine using the extensive and well-characterized ADNI dataset the hypothesis that ACC metabolism in healthy seniors functions as a mediator in the relationship between age and executive function. In agreement with our previous findings, highly significant correlations arose between age and metabolism; metabolism and fluency; and age and fluency. These observations motivated a mediation model in which ACC metabolism mediates the relationship between age and fluency score. Significance of the indirect effect was examined by Sobel testing and bootstrapping. In these cognitively intact seniors with “typical aging,” there was neither a correlation between age and memory scores nor between ACC metabolism and memory scores. The metabolism in a control region, the primary motor cortex, showed no correlation with age or ACC metabolism. These findings motivate further research into aging-related ACC dysfunction to prevent, diagnose, and treat the decline in executive function associated with aging in the absence of known neurodegenerative diseases.

**SIGNIFICANCE STATEMENT:** The pathophysiology of aging-related cognitive decline remains unclear but the anterior cingulate cortex (ACC), a major component of the anterior human attention system, shows decreasing metabolism that correlates with declining executive function despite otherwise intact cognition. Here, the relationships between ACC metabolism, age, executive function, and memory were examined using the large, public, ADNI database. Earlier findings were confirmed. In addition, ACC metabolism was found a mediator between age and executive function. In contrast, no correlation arose between memory and age or between memory and ACC metabolism. No correlations surfaced when using the metabolism of the right primary motor cortex as a control region. Development of preventive medicine and novel treatments will require elucidation of aging-related ACC pathophysiology requiring further research.

## INTRODUCTION

Cognitive aging (CA), also termed age-associated cognitive decline, describes the decrease in cognitive performance observed in otherwise healthy, cognitively intact individuals free of known neurodegenerative disorders. Three cardinal cognitive factors that decline are processing speed, memory, and executive function. In contrast, acquired skills, language, and knowledge about facts remain preserved relatively (Salthouse, 2010).

Much previous work in CA may have included individuals that are cognitively intact but not normal. The relationship between CA and neurodegenerative disease, if any, has thus remained unclear. Excluding subjects with preclinical Alzheimer’s disease (AD) or very early AD was impossible until recently—and remains difficult even today. AD is the most common dementing illness affecting one out of every three elders 85 years and older. Since AD begins 10-20 years before clinical presentation, a substantial fraction of seniors when enrolled in CA studies may have an evolving AD neurodegenerative process.

Exclusion of such individuals from CA research required traditionally prolonged follow-up with neuropsychological assessment. More recently, biochemical studies (e.g., amyloid, tau, phosphotau) of cerebrospinal fluid and as well as neuroimaging can identify individuals with preclinical (also known as asymptomatic) AD or early AD. Studies using positron emission tomography of glucose metabolism (^18^F-fluorodeoxyglucose; FDG PET), amyloid, and tau have become useful diagnostic adjuncts. Nevertheless, the specific biomarkers defining preclinical AD require further refinement. One approach considers the presence of fibrillar amyloid (A), tau neurofibrillary tangles (T), and neurodegeneration (N) (Sperling et al., 2011; Jack et al., 2018). However, the degree (i.e., threshold for positivity); kind (e.g., diffuse vs. dense-cored plaques); localization (i.e., region of interest); concordance with CSF markers; and potential new biomarkers are perforce an evolving definition of preclinical AD and its continuum.

The earliest neuroimaging biomarkers for AD are hypometabolism, atrophy, and amyloid deposition in the posterior cingulate cortex (PCC) and precuneus as well as in other parietal cortices (Minoshima et al., 1994; Rowe et al., 2007; Chételat et al., 2016). With PET imaging, tau deposition often follows in the inferior and medial temporal lobes leading to hypometabolism and cortical atrophy in those areas as well (Ossenkoppele et al., 2016; Scholl et al., 2016; Sintini et al., 2019). Of note, these neuroimaging findings can appear to diverge from those seen in some neuropathological studies (Braak and Tredici, 2015).

In contrast, human CA is not associated with PCC hypometabolism but affects instead predominantly the ACC and immediately adjacent regions (Pardo et al., 2007; Vaidya et al., 2007). The ACC is a central structure in human attention involved with many executive functions such as selective attention, resolution of conflict, multitasking, and adaptation to changing contexts (e.g., Pardo et al., 1991; for review, see Heilbronner and Hayden, 2016). The aging-related decline in ACC glucose metabolism correlates with the decline in executive function based on scores from a cognitive acuity screen as well as a commonly used test of executive function such as verbal category semantic fluency (Pardo et al., 2007). Separating effects of ACC hypometabolism on memory, as contrasted with effects on executive function, bears caution given the requirements for attention on memory (e.g., retrieval) and vice-a-versa (Norman, 1968; Downing, 2000; Chun and Turk-Browne, 2007).

Hypometabolism and cortical thinning can occur where amyloid deposits, but not necessarily (e.g., Altmann et al., 2015). In contrast, the later localization of tau deposits more closely correlates with regions showing hypometabolism or atrophy as well as clinical symptoms (e.g., Ossenkoppele et al., 2016). In AD, amyloid frequently involves also the ACC but not typically without incipient AD (Scholl et al., 2016).

The current study aims to examine the relationships between age; ACC metabolism; and cognitive functions such as memory and executive function. The study was motivated by the need to define pathophysiological mechanisms of aging-related phenomena within the ACC—a prerequisite to develop prevention strategies and treatments for the cognitive decline that occurs in aging uncontaminated by preclinical AD or early AD.

## MATERIALS AND METHODS

### Participants

Data from 231 participants aged 59-93 years (108 males, 123 females, mean age 74 years, SD 6) assessed as cognitively normal were obtained from LONI. Participants classified as cognitively normal had a score between 24-30 on the Mini-Mental State Exam (MMSE); had a Clinical Dementia Rating (CDR) of 0; and were not MCI, demented, or depressed (see ADNI for further diagnostic criteria). These participants had completed 6 or more years of education and were fluent in either English or Spanish. Participants’ ages and scores on category verbal fluency and paragraph memory were obtained from ADNI and were appropriately matched to the mean glucose uptake in an ACC region of interest (ROI). Fluency was scored using the “animal” category (i.e., name as many animals as possible in 1 minute) and served as a measure of executive performance. Memory was scored using the Wechsler Memory Scale-Revised Logical Memory II (delayed paragraph recall; Wechsler, 1987). The rationale for these measures will be forthcoming in the discussion.

Additionally, any cognitively-normal subjects who were identified as amyloid positive by ADNI were excluded to examine the potential role of amyloid deposition in the observed results. This provided a separate group of subjects (N = 155). Cortical-to-whole-cerebellum standardized uptake value ratios (SUVrs) were calculated for ROIs (frontal, temporal, parietal, anterior cingulate, posterior cingulate, and precuneus). A threshold of 1.1 defined amyloid positivity using as reference the value for whole cerebellum (Clark et al., 2011; Landau & Jagust, 2014; Joshi et al., 2015).

### Imaging

FDG PET scans from the Alzheimer’s Disease Neuroimaging Initiative (ADNI@LONI) were downloaded. These were stereotactically normalized to standard Talairach space (Talairach and Tournoux, 1988) via the <u>Neurostat</u> program (S. Minoshima, University of Utah). The FDG data were normalized through proportional scaling to whole brain glucose uptake (1000 counts). An ROI analysis (11 mm diameter sphere centered on x = +3 mm (right), y = +17 mm, z = +36 mm; i.e., the maximum from Pardo et al., 2007) for each scan was conducted. The PET imaging protocols are documented at the ADNI website (http://adni.loni.usc.edu/data-samples/pet/).

### Experimental Design and Statistical Analyses

Data were obtained from the Alzheimer’s Disease Neuroimaging Initiative (ADNI) database (adni.loni.usc.edu). ADNI was launched in 2003 by the National Institute on Aging (NIA), the National Institute of Biomedical Imaging and Bioengineering, the Food and Drug Administration (FDA), private pharmaceutical companies, and nonprofit organizations. At its core, ADNI is a longitudinal study whose main goal is to examine whether serial magnetic resonance imaging (MRI), positron emission tomography (PET), other biological markers, and clinical and neuropsychological assessments can be collectively used to measure the progression of mild cognitive impairment (MCI) and early AD. ADNI’s principal investigator is Michael Weiner, MD, VA Medical Center and University of California, San Francisco. For up-to date information, see “http://www.adni-info.org.”

Correlational analyses were performed between the following variables: age, ACC metabolism, animal fluency scores, and logical memory. To test whether ACC metabolism was significantly affecting the relationship between age and fluency scores, a mediation model was constructed with ACC metabolism as the mediator if three conditions were met: 1) age was correlated with ACC metabolism, 2) age was correlated with fluency scores, and 3) ACC metabolism was correlated with fluency scores. In the mediation analysis, age was regressed onto fluency scores (direct effect); then ACC metabolism as a predictor was added along with age in a second regression onto fluency scores (indirect effect). The total effect is the sum of the direct effect and indirect effect. The Sobel (1982) test assessed the significance of the difference between the direct effect and the total effect. A significant difference would indicate ACC metabolism was a mediator in the relationship between age and fluency scores. To increase rigor in testing for the significance of the mediating effects of ACC metabolism, a bootstrap (Preacher and Hayes, 2008) with 5,000 simulations was performed to estimate 95% confidence intervals for mediation effects. All pathways were tested simultaneously. Analyses and computations were performed using R, version 3.5.1 (http://www.R-project.org/). The mediating or regression effect was considered significant (i.e., different from 0) at the corresponding confidence level if the upper and lower bounds of the confidence interval had the same sign.

## Results

### Mediation Analysis: Relationships between age, ACC metabolism, fluency, and metabolism in a control region in cognitively intact seniors

Age was a significant predictor of fluency (c = −0.24, SE = 0.06; p = 1.7(10)^−4^; Fig. 1A). Age was also a significant predictor of ACC metabolism (a = −0.44, SE = 0.06, p = 2.6(10)^−12^; Fig. 1B). Age along with ACC metabolism was a significant predictor of fluency scores (b = 0.15, SE = 0.07, p = 0.03; Fig. 1C). Controlling for ACC metabolism (mediator), the predictive effect of age on fluency scores, although still significant, decreased (c′ = −0.18, SE = 0.07, p = 0.01; Fig. 1D) consistent with partial mediation. The Sobel test indicated that the reduction in the effect of age on fluency after controlling for ACC metabolism (indirect effect) was significant further showing the mediation effect of ACC metabolism was significant (Sobel Statistic = −2.03, p = 0.04). Approximately 19% of the variance in fluency scores was accounted for by the predictors (R^2^ = 0.193). A bootstrap estimation with 5,000 simulations tested the significance of the indirect effect or the mediating effect (Shrout and Bolger, 2002). The results indicated the mediating effect of ACC metabolism approached significance (β = −0.06, 95% CI = −0.14160, 0.00, p = 0.059). Like the results of the regression analysis, the bootstrap estimation showed a decrease in the effect of age on fluency when controlling for ACC metabolism (decrease in absolute value between Total Effect and ADE, Table 2). Additionally, there was a significant correlation between fluency and memory scores (Table 1).

**Figure 1.**
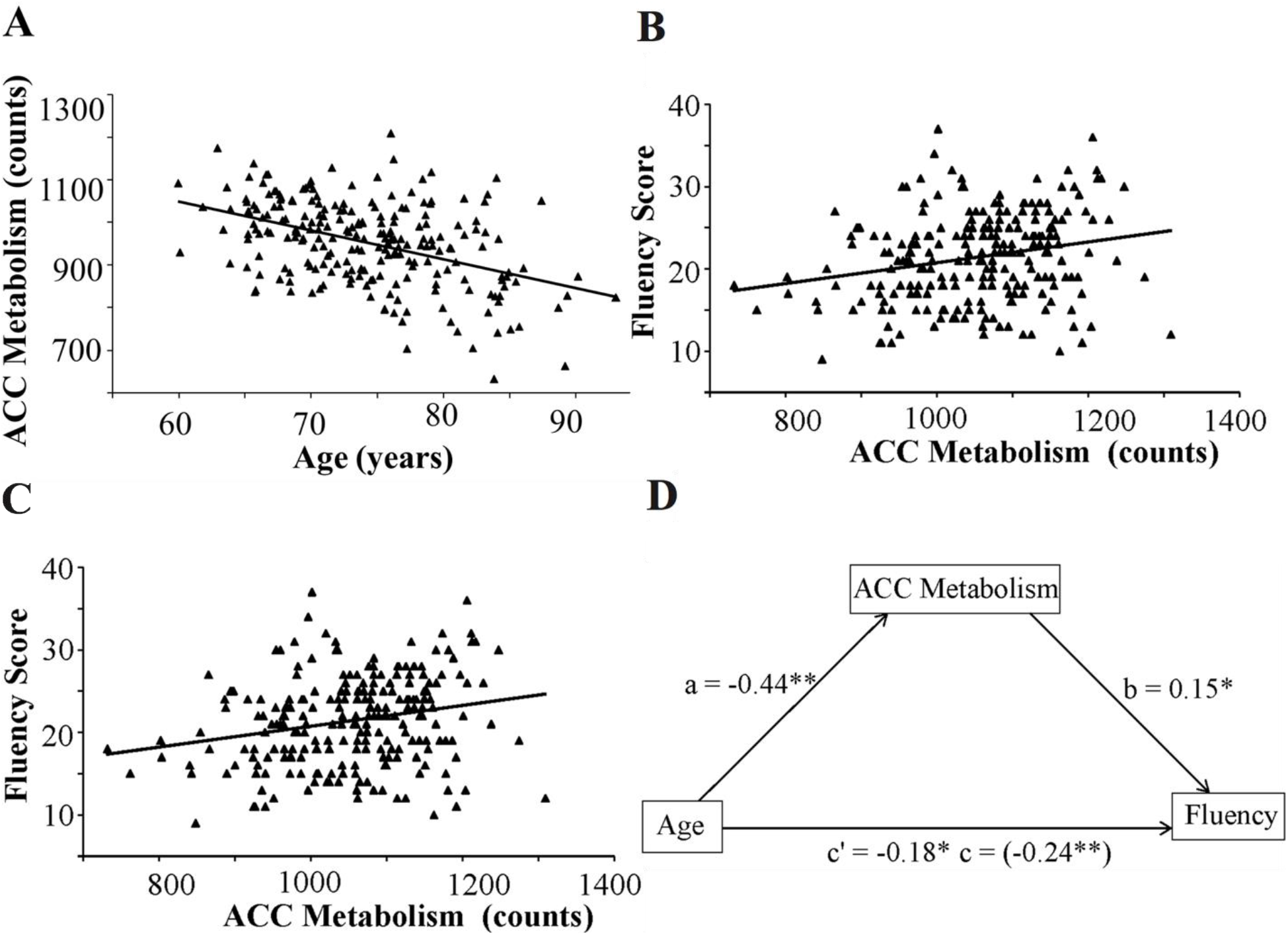
Mediation model with ACC metabolism as a mediator in the relationship between age and fluency. Value in parenthesis represents total effect. * p < 0.05. **p < 0.001. **A.** Correlation between age and ACC metabolism replicating prior results with ADNI dataset (Pardo et al., 2007). **B.** Correlation between age and fluency. **C.** Correlation between ACC metabolism and fluency. **D.** ACC metabolism is a partial mediator of the relationship between age and fluency.

**Table 1.** Correlations between age, ACC metabolism, fluency scores, and memory scores.

|  | Age | ACC Metabolism | Fluency | Memory |
| --- | --- | --- | --- | --- |
| Age | 1.00 | -0.44** | -0.25** | 0.08 |
| ACC Metabolism | — | 1.00 | 0.23** | -0.01 |
| Fluency | — | — | 1.00 | 0.18** |
| Memory | — | — | — | 1.00 |

\* $p \leq 0.05$
\*\* $p \leq 0.01$

**Table 2.** Nonparametric bootstrap: ACC metabolism as a mediator of age and fluency.

|  | **Estimate** | **95% CI** | **p-value** |
| --- | --- | --- | --- |
| **ACME ($\beta$)** | -0.06632 | [-0.14160, 0.00] | 0.059 |
| **ADE** | -0.17854 | [-0.31452, -0.04] | 0.012* |
| **Total Effect** | -0.24487 | [-0.35848, -0.13] | $<2(10)^{-16}$** |
| **Prop. Mediated** | 0.27085 | [-0.00926, 0.74] | 0.059 |

Simulations = 5,000
ACME = Average Causal Mediation Effects
ADE = Average Direct Effects
\* $p \leq 0.05$ ; \*\* $p \leq 0.01$

A similar analysis determined whether metabolism in the right motor cortex (RMC) mediates the effect of age on executive function. Contrary to the results seen for ACC metabolism as a mediator, age was not a significant predictor of RMC metabolism (a = −0.04, SE = 0.07, p = 0.51; Fig. 2); neither was RMC metabolism a significant predictor of fluency scores (b = −0.05, SE = 0.06, p = 0.45; Fig. 2). When RMC metabolism was controlled, the predictive effect of age on fluency scores decreased by a small amount but remained significant (c′ = −0.25, SE = 0.06, p = 1.5(10)^−4^). The Sobel test showed that this reduction was not significant (Sobel Statistic = 0.50; p = 0.62). A bootstrap estimation with 5,000 simulations to test the mediating effect also indicated that the mediating effect was not significant (β = 0.002, 95% CI = −0.007, 0.01, p = 0.73). The bootstrap estimation did not show a decrease in the effect of age on fluency when controlling for RMC metabolism (not shown).

**Figure 2.**
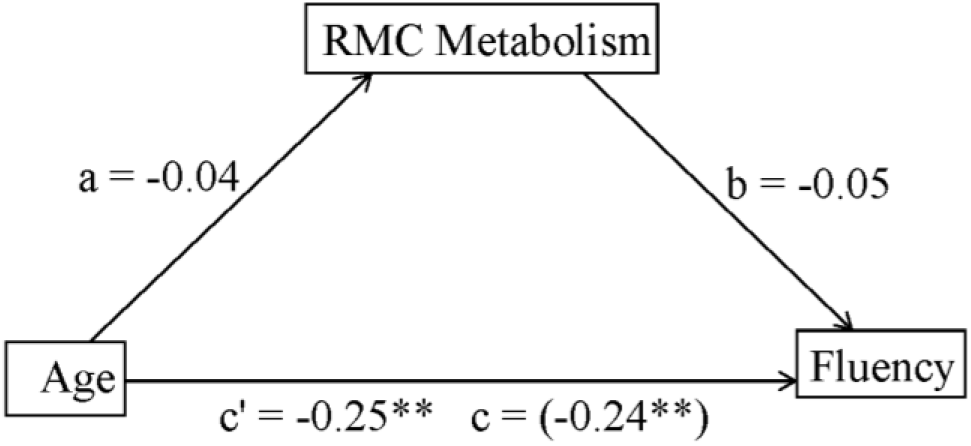
Mediation model with primary motor cortex metabolism as a mediator in the relationship between age and fluency. Value in parenthesis represents total effect. *p < 0.05. **p < 0.01.

### Associations with memory in cognitively intact seniors

No significant correlation was observed between logical memory scores and age (r = 0.08, p = 0.19; Fig. 3A) or ACC metabolism (r = −0.01, p = 0.92; Fig. 3B). Given the lack of a significant correlation, no mediation analysis was warranted. With regards to the control region, i.e., the right primary motor cortex (RMC), no significant correlation was observed between RMC metabolism and either age (r = −0.04, p = 0.51) or logical memory (r = −0.004, p = 0.96).

**Figure 3.**
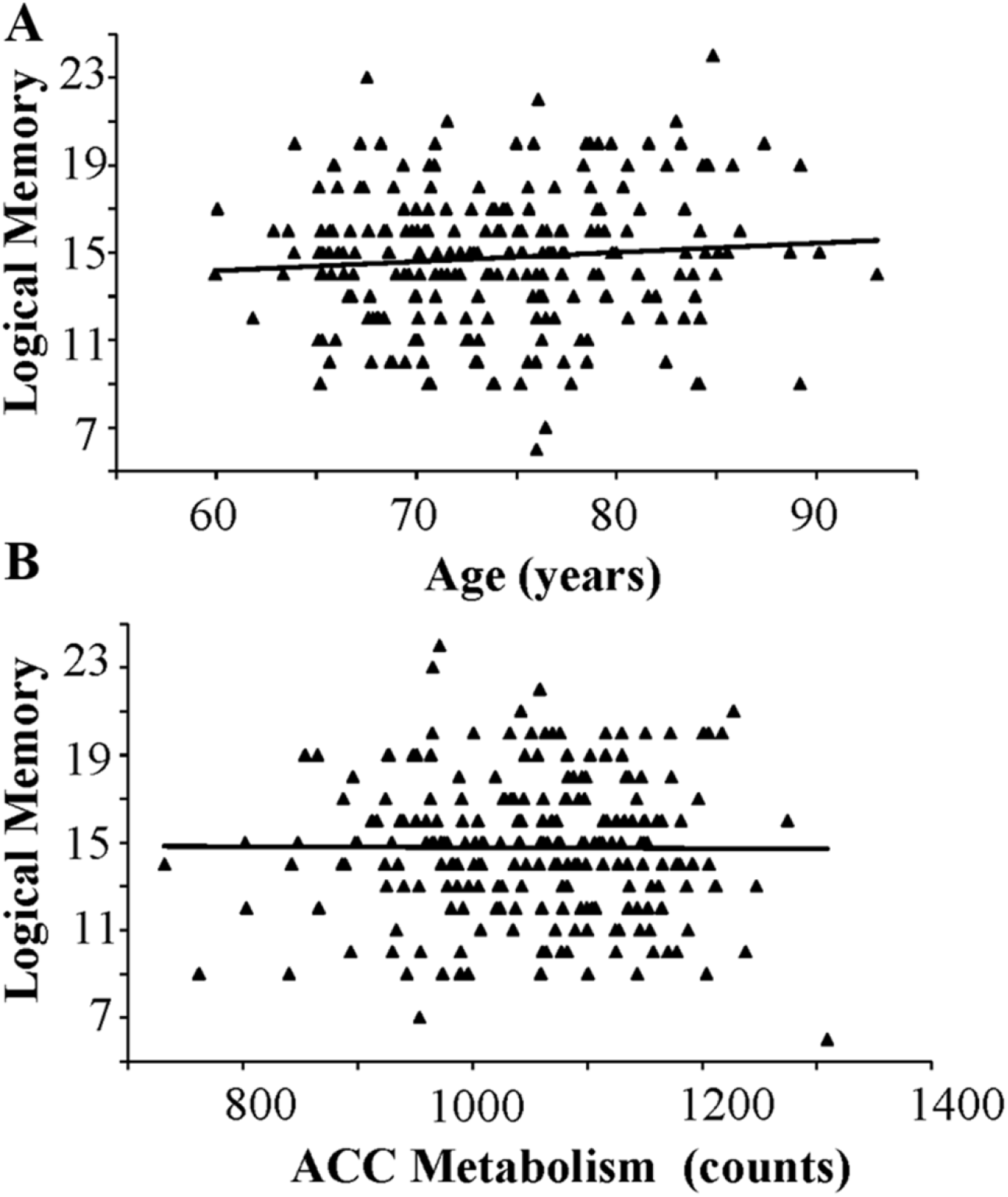
Correlation between scores on the test of logical memory and either age (A) or ACC metabolism (B).

### Effects of amyloid positivity

As discussed previously, contamination of cognitively normal subjects by individuals with preclinical or very early AD can confound the study of CA. In the current state of knowledge, it is admittedly not possible to completely rule out preclinical AD. Nevertheless, the previous analyses were repeated using only amyloid negative, cognitively normal subjects from ADNI that included 155 subjects. This group had at least a lower likelihood of preclinical AD. ADNI’s designation of amyloid positivity was SUVR 1.1 derived from critical ROIs.

ACC metabolism became a complete mediator of the relationship between age and fluency as the direct effect (c′) was no longer significant (as in Fig. 1: a = −0.51, SE = 0.07, p = 8.8(10)^−12^; b = 0.22, SE = 0.09, p = 0.01; c = −0.27, SE = 0.08, p = 7.8(10)^−4^; c′ = −0.15, SE = 0.09, p = 0.09). The Sobel test confirmed the mediation (Sobel Statistic = −2.34, p = 0.02). In comparison to the previous bootstrap analysis showing a trend level of significance (Table 2, β), the reanalysis using only amyloid negative subjects reached significance confirming ACC metabolism mediated the relationship between age and fluency (as in Table 3: β = −0.11, 95% CI = −0.2193, −0.02, p = 0.02). The reanalysis demonstrated once again no relationship between age and memory or between ACC metabolism and memory (r = 0.10, p = 0.12; r = 0.09, p = 0.16 respectively). In examination of the relationship between the metabolism in a control region (i.e., right motor cortex), age, fluency, and memory (i.e., analogous to Fig. 2), the only significant relationships were between age and fluency (as above) and between memory and fluency (r = 0.19, p ≤ 0.01). As expected, there was no significant correlation with the metabolism in the motor cortex.

**Table 3.** Nonparametric bootstrap: ACC metabolism as a mediator of age and fluency (amyloid negative subjects, N = 155).

|  | Estimate | 95% CI | p-value |
| --- | --- | --- | --- |
| **ACME ($\beta$)** | -0.1129 | [-0.2193, -0.02] | 0.0204* |
| **ADE** | -0.1541 | [-0.3218, 0.02] | 0.0888 |
| **Total Effect** | -0.2670 | [-0.3977, -0.13] | $<4(10)^{-4**}$ |
| **Prop. Mediated** | 0.4229 | [0.0599, 1.16] | 0.0208* |

Simulations = 5,000
ACME = Average Causal Mediation Effects
ADE = Average Direct Effects
\* $p \leq 0.05$ ; \*\* $p \leq 0.01$

## DISCUSSION

These results indicate ACC metabolism mediates the relationship between aging and executive function implicating some unknown pathophysiology in aging-related cognitive decline. In contrast, mnemonic dysfunction, a cardinal feature of AD, could not be implicated in the relationship between CA and ACC metabolism. The metabolic effects of age were specific to ACC as no relationship surfaced when examining the RMC indicating some regional specificity to the effects of age on metabolism.

### ACC metabolism: Mediator between age and fluency in healthy older adults

Previous results that spanned a broad range of ages (18-90 years) with a more limited sample size of 46 subjects reported a decline in ACC metabolism that correlated positively with declining executive function as measured by the same animal fluency task (Pardo et al., 2007). The decline in ACC flow or metabolism with age had been reported previously by several groups; however, the relationship of ACC metabolism to executive function in CA had not been noted previously. The subset of the ADNI dataset used here has a much larger sample size (N = 231) but narrower age range (56-92 years) and does not include young adulthood. The ADNI data enabled enough power to show that ACC metabolism was a mediator of the relationship between age and fluency in more elderly, cognitively normal adults. These results support the idea that ACC dysfunction plays a crucial role in age-associated cognitive decline (AACD). As such, ACC hypometabolism could serve as a potential biomarker for CA.

### ACC metabolism: Not a mediato between age and memory in healthy older adults

Episodic memory is often tested using lists of words or recall of items from a story presented at an earlier time. Recall of story items takes advantage of semantic processes, while recall of word lists requires self-generation of mnemonic strategies. Memory of word lists taxes the executive system more than does story recall. For example, patients with frontal lesions and those with executive dysfunction do worse on recall of word lists than on recall of items from paragraphs presented previously (Kopelman and Stanhope, 1998; Brooks, Weaver, and Scialfa, 2006). In contrast, memory for semantically-related information depends more on medial temporal processes (Tremont et al., 2000; Lezak, Howieson, and Loring, 2004). Therefore, when attempting to differentiate the ACC’s role in executive vs. mnemonic processes, the Logical Memory test is preferred. ACC metabolism does not mediate the relationship between age and Logical Memory. In fact, these cognitively intact subjects did not show significant correlations between age and Logical Memory or between memory scores and ACC metabolism. Of interest, there was a significant correlation between memory and fluency scores highlighting interdependence between memory and attention despite the very large correlation carried by age and metabolism.

### Limitations

#### ROI definition and multiple comparisons

This research was motivated specifically by our prior results of ACC metabolism in CA (Pardo et al., 2007). As such, an exhaustive search for other areas that might show a high correlation with age was not done. In part, problems in ROI definition and concerns about multiple comparisons tempered such an approach.

#### Preclinical AD

Another issue is concern about possible inclusion of preclinical AD in the cognitively intact elderly sample. The absence of memory changes with age in the cognitively intact seniors suggests minimal contamination with subjects having preclinical AD or early AD. However, complete exclusion of preclinical AD remains problematic even today.

The exclusion of amyloid positive subjects made the results more robust. ACC metabolism moved from a partial to a total mediator of the relationship between age and fluency.

A few individuals converted to AD shortly after the baseline evaluation identified them as normal—for example, elderly APOE4 homozygotes (Pardo and Lee, 2018). Curiously, these elderly, cognitively intact APOE4 homozygotes did not show significant PCC hypometabolism or PCC amyloid deposition. Yet, the PCC shows the earliest signs of hypometabolism in AD (Minoshima et al., 1994). It remains debated whether a patient with dementia and without any amyloid deposition (diffuse or fibrillar) can be diagnosed with AD. For example, Hyman et al. (2013) in an NIA/Alzheimer’s Association consensus conference note “…a major point of discussion among committee members was the relative value of evaluating both Ab/amyloid plaque phase and neuritic plaque score…in the assessment of AD neuropathologic change. Because the relative independent value of these two parameters is not currently known, we suggest collecting data on both and evaluating their independent value in future analyses.

Cross-sectional neuropathological studies suggest that a significant number of demented individuals with extensive tau pathology have minimal fibrillar plaques (Braak & Tredici, 2015; Monsell et al., 2015). Amyloid PET cannot detect diffuse plaques; so, this remains a possibility for those with negative amyloid scans. Even amnestic, amyloid-negative patients (mostly without APOE4) initially diagnosed as AD, whose clinical phenotype remained AD over a follow-up interval of four years, showed PCC hypometabolism and atrophy (Chételat et al., 2016). Of note, a negative amyloid scan in the IDEAS study prompted changing management in 2/3 of patients within 90 days of the scan. Ultimately, the precise criteria for exclusion of preclinical AD will need refinement: CSF criteria, pretangles, neurofibrillary tangles, diffuse plaques, neuritic plaques, types and extent of neurodegeneration, and probably new forthcoming biomarkers.

#### Atrophy and partial volume correction

The PET data were not corrected for partial volume effects (PVC). ACC hypometabolism could reflect focal atrophy without a change in grey matter metabolism. However, PVC can introduce noise especially when volume changes are small. Vaidya et al. (2007) reported that even with PVC, the decline in ACC blood flow (correlated with metabolism) with age could not be totally accounted for by atrophy. The correlations between flow and age were much larger than correlations between age and grey matter volume, and some regions showed strong correlation with age without a change in grey matter volume.

Whether the ACC shows any atrophy with age remains controversial. Two large cross-sectional studies using scans at high resolution found thickening of the ACC grey matter with aging (Salat et al., 2004; Fjell et al., 2009). However, a longitudinal study also with a large sample size and high-resolution structural scans did not find ACC thickening with age over a three-year period, but neither did it show gross atrophy (Fjell et al., 2014). Another longitudinal study of cognitively intact elders did not show significant ACC volume changes over 4 years but did find significant decreases in the ACC glucose uptake even with correction for partial volume effects (Catellanos et al., 2019).

There is also evidence ACC thickness may be a trait or state marker. Aged individuals who have memory performance akin to those who are much younger (i.e., “SuperAgers”) show a thickened ACC (Rogalski et al., Geffen et al.) Education serves often as a proxy for cognitive reserve. In amyloid-negative, healthy elders the years of education correlate positively with ACC volume, ACC metabolism, and ACC functional connectivity with the hippocampus and PCC (Arenaza-Urquijo et al., 2013). Furthermore, sedentary elders show increased ACC volume in a between-group comparison following an intervention with aerobic vs. stretching execises (Colcombe et al., 2009).

#### Correlation in cross-sectional studies

Correlational and cross-sectional studies can confound interpretation. Different pathophysiologies may participate in ACC dysfunction during different time intervals in the aging brain. For example, most loss in AG occurs before age 55 years (Goyal et al., 2017). In contrast, decreased glucose uptake may occur later in life when peripheral insulin resistance arises (Castellano et al., 2019).

#### Association vs. causality

Association does not prove causality. Regional ACC effects could come from extrinsic regions; i.e., ACC dysfunction is an effect rather than a cause. However, no other brain region showed greater decline in metabolism with age. Nevertheless, the results highlight the ACC as a critical region to look for pathophysiology which will require an animal model.

Preclinical models remain unsettled whether aging rodents show a correlation of mPFC hypometabolism with declining executive function. The human ACC/mPFC homologues in rodents are under refinement (Vogt and Paxinos 2014). Furthermore, optimal probes of rodent executive function are unclear.

Several paradigms probe executive function in rodents. Older rats perform poorly on the water maze task and show, among other regions, decreased ACC metabolism (Gage et al., 1984). Representation of 3D-objects in remote memory recruits the ACC (Burke and Barnes, 2015). Ibotenic acid lesions targeting the PL ACC selectively impaired performance on extradimensional set shifting (Birrell et al., 2000). Chemogenetic inhibition of the mouse dorsal ACC impaired sustained attention as indexed by a 5-choice serial reaction time task (Koike et al., 2016). Selective lesions of different medial prefrontal regions (Cg, PL) affected preferentially different components of set shifting and probabilistic reversal learning (Ng et al., 2007; Newman and McGaughy, 2011; Dalton et al., 2016).

In summary, ACC metabolism mediates the relationship between age and executive function while having no relationship to memory in cognitively intact elders. This relationship does not occur throughout the brain consistent with the known specialization of the ACC for executive functions. These findings motivate research on aging-related ACC pathophysiology and the need for animal models of CA.

## Acknowledgments

Grant Funders: Department of Veterans Affairs (CSRD) # I01CX000501 (JVP) and the Minneapolis Veterans Health Care System. The authors thank the participants of ADNI as well as the ADNI researchers for making this resource open to the public.

